# The absence of the Type VI Secretion System in the successful lineage of *Acinetobacter baumannii* ST19

**DOI:** 10.1101/2025.04.29.651177

**Authors:** Adam Valcek, Diana Isabela Costescu Strachinaru, Kristina Nesporova, Patrick Soentjens, Anke Stoefs, Charles Van der Henst

## Abstract

*Acinetobacter baumannii* is an opportunistic pathogen, often multi- to pandrug-resistant, including to last-resort antibiotics such as carbapenems. *A. baumannii* can acquire DNA through multiple mechanisms including natural competence and conjugation. An active Type VI Secretion System (T6SS), which kills nonkin bacteria and challenges and inhibits the conjugation. The acquisition of resistance genes has helped the spread of resistant clones of *A. baumannii*, such as those of Sequence Types (ST^Pas^), ST1, ST2, ST10, ST15, ST25, ST79 and ST85. While ST1 and ST2 were thoroughly studied, there are new emerging clones whose genetic background facilitating the rise is still unknown or poorly understood.

The emergence of *Acinetobacter baumannii* ST19 first peaked in 2003, coinciding with the Iraq conflict and increased detection among U.S. military personnel. Its prevalence later surged in Georgia and Ukraine, around the onset of the conflict in Ukraine in 2022, associating it with regions affected by military activity.

In this study, we have sequenced whole-genome three isolates of *A. baumannii* ST19 and compared them with further 156 *A. baumannii* ST19 genomes were obtained from public repositories. In total, 157/159 genomes revealed loss of T6SS locus, which was replaced in 56/157 genomes (35,7%) by ΔTn*9* carrying chloramphenicol and ΔTn*10* carrying tetracycline resistance genes, and formaldehyde and chlorite resistance genes. Surprisingly, the antibiotic resistance-encoding transposons likely originated from Enterobacteriaceae plasmids. The loss of a functional T6SS in *A. baumannii* ST19 may potentially facilitate horizontal gene transfer and promoting a cooperative or less competitive lifestyle providing a selective advantage at the population level.

## Introduction

*Acinetobacter baumannii* is an emerging opportunistic nosocomial pathogen that causes a variety of infections including urinary tract infections, wound infections, meningitis, ventilator-associated pneumonia, blood-stream infection, and endocarditis (1). *A. baumannii* is a member of ESKAPE pathogens (*Enterococcus faecium*, *Staphylococcus aureus*, *Klebsiella pneumoniae*, *Acinetobacter baumannii*, *Pseudomonas aeruginosa* and *Enterobacter* spp.) (2), and due to its frequent resistance to last-line antibiotics such as carbapenems, it has been designated as a critical priority in the list of pathogens of importance for studying and developing new antibiotics by the World Health Organization’s list (3, 4).

Besides conventional antibiotics, *A. baumannii* can withstand challenging environmental conditions such as oxidative stress, prolonged periods of desiccation, human serum but also disinfectants (5).

The ability of *A. baumannii* to acquire exogenous DNA via natural competence (6) and conjugation (7) contributes to high plasticity of its genome (6). The horizontal gene transfer plays significant role in the evolution of *A. baumannii* lineages, resulting in number of highly successful clonal lineages such as ST1, ST2, ST10, ST15, ST25, ST79 and ST85 (Pasteur scheme) (8, 9). While ST1 and ST2 are responsible for the majority of infections caused by extensively-drug resistant *A. baumannii*, some clones are specific to certain geographical locations (9), such as ST25 (Europe, United Arab Emirates, South America, Asia) (10), ST32 (South Korea, Europe, the USA, Canada and the Middle East) (11) or ST78 (Italy, Ukraine, USA) (12).

While the horizontal gene transfer, especially plasmid transfer via conjugation, can shape entire lineages and their success (13), an active Type VI secretion system (T6SS) poses a unique challenge to conjugative plasmids, as *Acinetobacter* plasmid donors and recipients may kill each other (14). The T6SS machinery is usually encoded in a single chromosomal locus (15). The T6SS of *A. baumannii* has been shown to mediate the killing of *E. coli*, *K. pneumoniae* and *P. aeruginosa*, while different strains of *A. baumannii* are also able to kill each other (15, 16).

In this study, we have assessed genetic determinants disrupting the T6SS-encoding locus in a rising lineage of *A. baumannii* ST19, of which occurrence is increasing regionally (Ukraine, Georgia) (17). Furthermore, we have performed comparative genomic analysis of the global population of *A. baumannii* ST19 in order to decipher genomic traits that make this lineage emergingly successful and to assess its threat in becoming a world-wide disseminated clone.

## Results

### *A. baumannii* ST19 strains and genomes

A concerning report on the increasing occurrence of *Acinetobacter baumannii* ST19 encoding carbapenemases was recently published (17), raising questions about factors contributing the success of this lineage. Three isolates of *A. baumannii* ST19 were obtained from two different patients (patient 1 - wound, rectal swab; patient 2 - rectal swab) and long-read whole-genome sequenced and *de novo* assembled.

A further 156 genomes (including 97 from Luo et al. (17)) of *A. baumannii* ST19 were retrieved from public repositories (GenBank and BIGSdb). The occurrence of *A. baumannii* ST19 appears to spike during specific time periods that correlate with military conflicts (Figure 1), such as the Iraq War (2003) and the war in Ukraine (2022).

**Figure 1:**
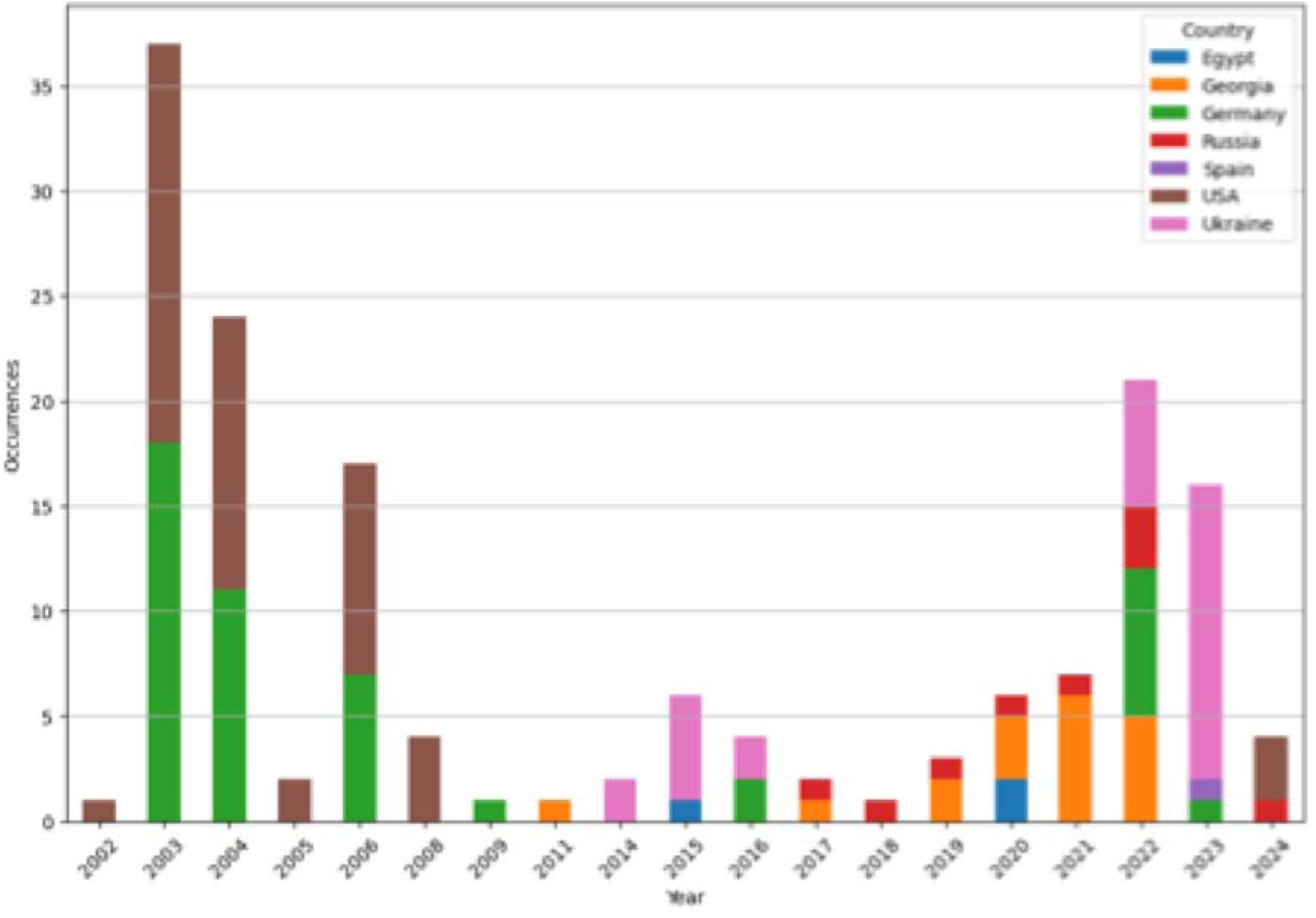
Temporal and spatial occurrence of *A. baumannii* ST19 based on genomes deposited in public repositories and available metadata.

The examined genomes carried a variety of virulence and resistance genes (Supplementary Figure 1), including resistance to last-resort antibiotics such as carbapenems. One-third (n=53) out of 159 genomes described in this study encoded at least one carbapenemase gene (*bla*_OXA-23_, *bla*_OXA-72_, *bla*_NDM-1_ or *bla*_NDM-5_) with 6 encoding two and 9 encoding three carbapenemases. Furthermore, a broad variety of plasmid replicons was detected (Supplementary Figure 1), suggesting an easy acquisition of genetic material by the horizontal gene transfer.

Genomic analysis suggests that the possible underlying reason for its success to be the acquisition of multiple partial transposons, including ΔTn*9* (18) carrying chloramphenicol resistance gene and ΔTn*10* (19) carrying the tetracycline resistance gene. Interestingly, 61/159 (38,4 %) genomes of *A. baumannii* ST19 carried chlorite dismutase and formaldehyde resistance (*frmRAB*) genes, with 98,4% amino acid identity to *Massilia glaciei* and 96,7% to *Ideonella dechloratans*, where it combats oxidative stress (20). To our knowledge, this is the first time chlorite dismutase has been reported in the genome of *A. baumannii*.

Furthermore, these transposons disrupted and replaced the locus encoding the T6SS. The entire region of T6SS deletion exhibits extensive genetic rearrangements, as ΔTn*9* and ΔTn*10* were acquired and disrupted by IS*26* and IS*4321,* and IS*26*, respectively.

The absence of a functional T6SS, likely enabling conjugal acquisition of genetic material (14) such as antibiotic, formaldehyde and chlorite resistance, could represent one of the reasons behind their successful rise.

### Genetic variability of the *A. baumannii* ST19 lineage

The whole genome sequences were subjected to genotyping [sequence type, capsule locus (KL) type, outer core lipooligosaccharide locus (OCL) type, antimicrobial resistance, and virulence genes]. A phylogenetic analysis was performed on these isolates, including genomes of the same ST from public repositories (GenBank and BIGSdb) (Figure 2, Supplementary Figure 1).

**Figure 2:**
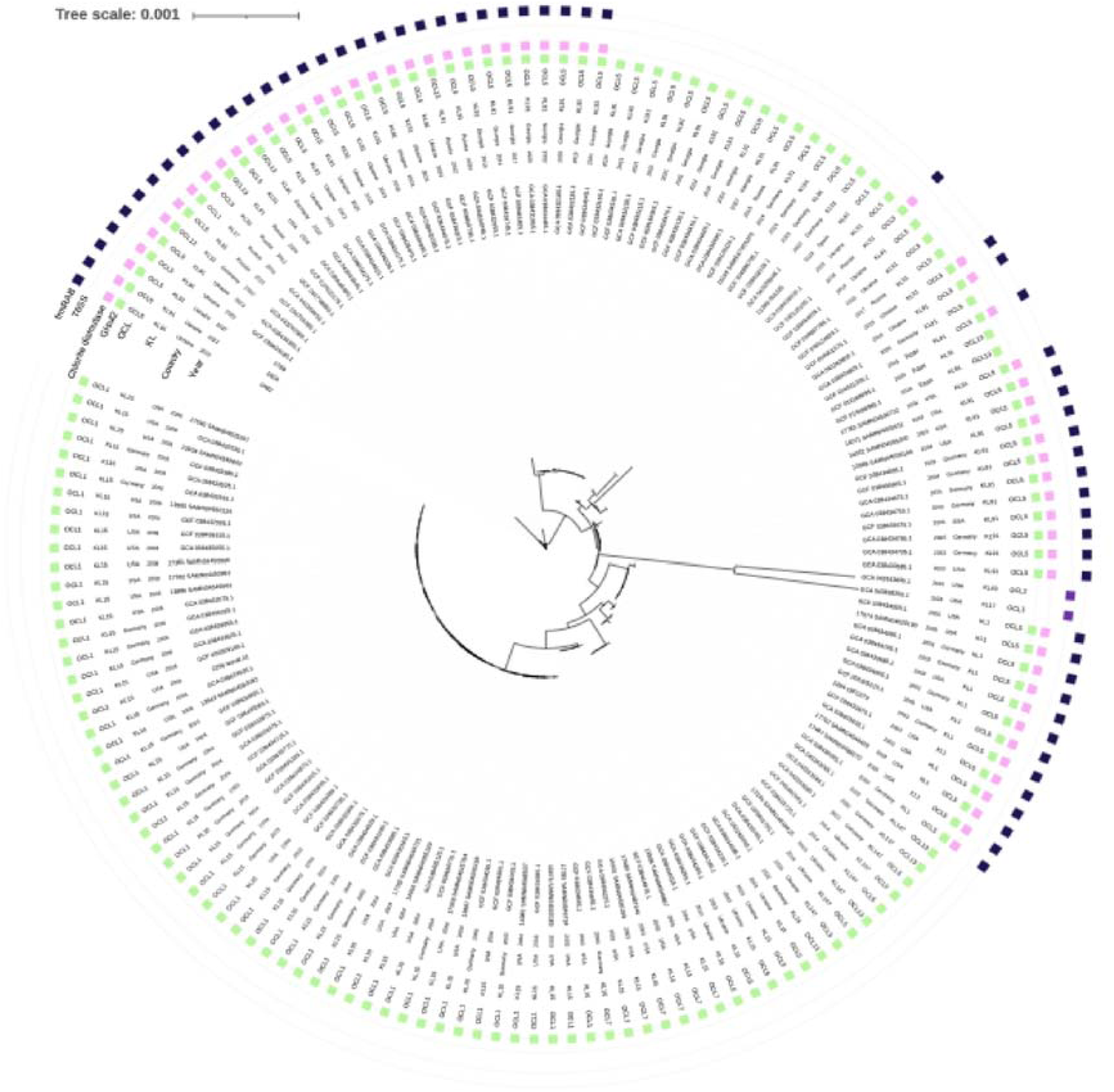
Phylogenetic analysis of 3 isolates and further 156 of *A. baumannii* ST19 genomes, depicting year of isolation, country, capsule locus type (KL), outer core lipooligosaccharide type (OCL), and presence of GI*sul2* genetic island, chlorite dismutase, T6SS, *frmRAB* genes, respectively.

While various MLST (Multilocus Sequence Type) ST (Pasteur scheme) such as ST2 or ST1 often belong to various KL types (6), the ST19 lineage exhibited limited variability with 69/159 genomes being KL91, 66/159 KL15, followed by KL1 (13/159), KL147 (8/159), KL17 (2/159) and KL40 (1/159). Interestingly, 53/55 OCL1 encoded KL15 (2/55 OCL1 were KL17), the most predominant OCL type, OCL5 (n=87) was variable and encompassed KL91, KL1, KL147 and KL15. Furthermore, all of the genomes of KL15/OCL1 formed one large, genetically related cluster (Figure 2).

The vast majority (157/159) of *A. baumannii* ST19 harbored GI*sul2,* a chromosomally carried <u>G</u>enomic Island encoding sulphonamide resistance determinant (*sul2*) along with putative arsenic resistance genes (21). We have detected a range of 0 to 6 plasmid replicons (22) in the genomes of *A. baumannii* ST19, with rP-T1 replicon being the most prevalent (n=122/159), followed by r3-T23 (n=46/159), r3-T12 and r1-T8 (n=44/159 both).

Interestingly, 18 isolates did not carry any detectable plasmid replicon raising a question whether there is another mechanism limiting acquisition of plasmids or whether they have not encountered a conjugative plasmid yet. However, the *Acinetobacter* typing schemes are still being developed (22), possibly undetected (or below the detection threshold) plasmids must be considered as well.

By evaluating the T6SS locus in the ST19 lineage of *A. baumannii*, we identified variability in genetic origin of the deletion of these genes. Of 61/159 genomes carrying chlorite dismutase and formaldehyde resistance genes in the locus of T6SS, 55/61 harbored ΔTn*10* carrying the tetracycline and ΔTn*9* carrying chloramphenicol resistance genes (Figure 3). Five formaldehyde resistance and chlorite dismutase-encoding genomes (5/61) did not harbor ΔTn*10* carrying the tetracycline resistance and 1/61 did not harbor ΔTn*9* carrying chloramphenicol resistance genes. However, mapping on the reference genome (0604 - SAMN46706488) confirmed, that in all these genomes (61/159) these genetic rearrangements displaced locus encoding T6SS.

**Figure 3:**
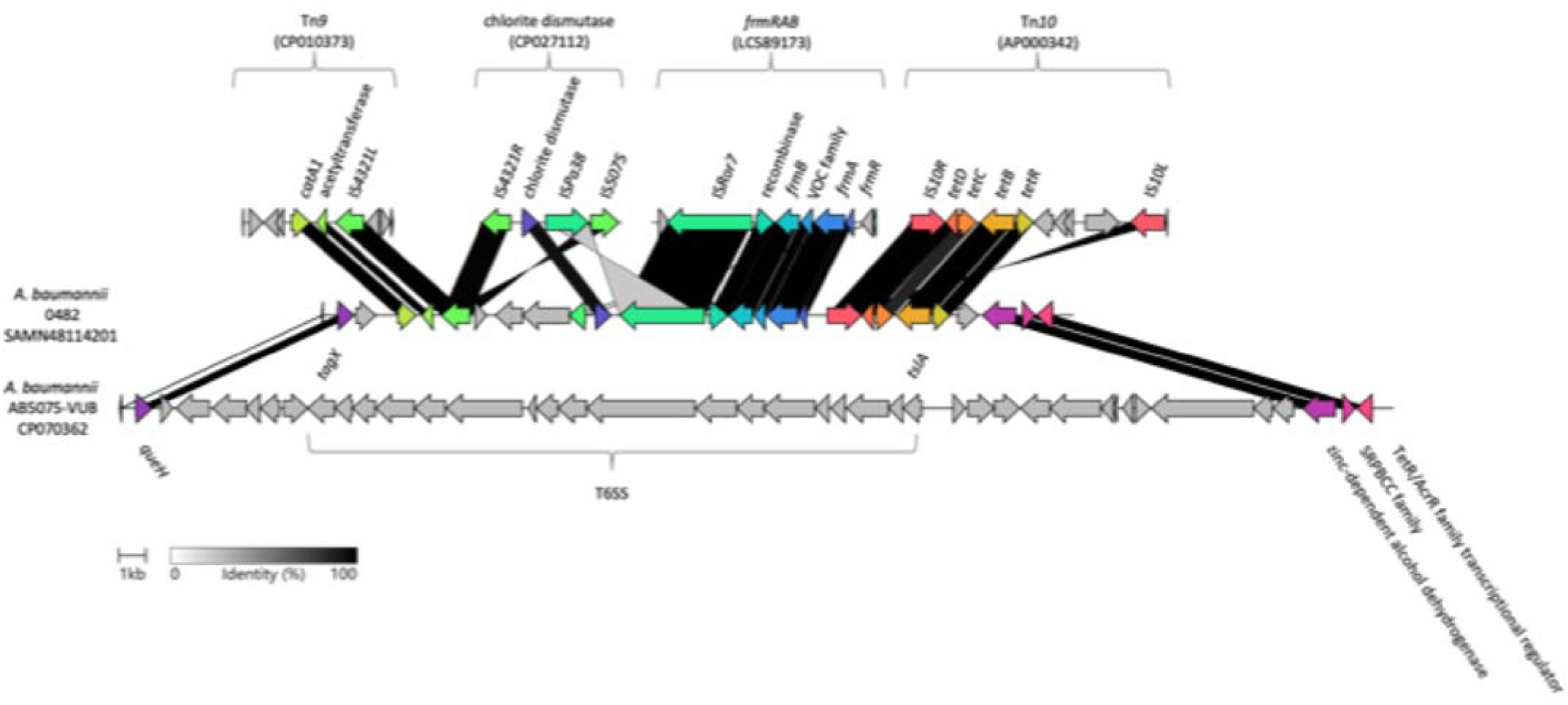
Depiction of the T6SS locus disruption of *A. baumannii* ST19 (0482 - SAMN48114201) as a representative genome, compared to AB5075-VUB (CP070362) showing deletion mediated by ΔTn*9*, chlorite dismutase, formaldehyde resistance locus *frmRAB*, and ΔTn*10*. The alignments are based on amino acid identity.

Interestingly, 2/159 genomes of *A. baumannii* ST19 (GCA_042643835.1 and GCA_040868255.1) still encoded a likely functional T6SS. However, compared to AB5075-VUB (accession nr. CP070362), a number of SNPs (99% pairwise identity over 19417 bp) was detected in these two genomes, while the broader genetic neighbourhood only sporadically exhibited SNPs. The genomes GCA_042643835.1 and GCA_040868255.1 belonged to KL40/OCL2 and KL17/OCL1, respectively, which are rare for this lineage (a sole KL40 and one of two KL17, respectively). While none of the SNPs were non-sense, the influence of missense or synonymous (e.g., via tRNA) on the functionality of T6SS remains to be explored.

### The loss of the T6SS in non-ST19 lineages of *A. baumannii*

The presence of the T6SS locus was assessed in 8218 genomes of *A. baumannii* available in BIGSdb. The lack of the T6SS was detected in 1287 genomes (including 159 genomes of *A. baumannii* ST19). The most dominant lineage of *A. baumannii* lacking the T6SS in this comparison was ST2 (n=435; 33,7%) followed by ST19 (n=159; 12,3%) and ST94 (n=82; 6,4%) (Supplementary Figure 3AB).

The phylogenetic comparison of all 1287 (Supplementary Figure 2) revealed clustering of all *A. baumannii* ST19 and 103 non-ST19 genomes together. The non-ST19 genomes in the ST19 cluster consisted of 93 single-locus variant (SLV) sequence types including one novel ST, and 10 double-locus variant (DLV) sequence types including one novel ST (Supplementary Figure 3C).

The presence of *frmRAB*, and chlorite dismutase-encoding genes was detected predominantly in *A. baumannii* ST19, but also in two *A. baumannii* ST315 genomes within the same cluster. A single genome (ST2525) outside of the ST19 cluster was found to carry the *frmRAB* genes.

## Discussion

*Acinetobacter* has the core components of the T6SS conserved and found in a single chromosomal locus (15). The *A. baumannii* strains use Type VI Secretion System (T6SS) to kill nonkin *A. baumannii* as well as other bacterial species such as *E. coli*, *K. pneumoniae* and *P. aeruginosa*, (16, 23). The active T6SS can be a challenge to conjugation and horizontal gene transmission (15). The absence of T6SS could therefore allow co-existence of nonkin bacterial species and strains. We hypothesize that this might and explain original acquisition of genetic material from these species and possibly evolution of the *A. baumannii* ST19 lineage.

The Tn*10* was originally identified and described in *Shigella flexneri* (19). This corresponds to the best match of the ΔTn*10* nucleotide sequence in this study was detected for *Salmonella. enterica* serovar Typhi and *S. flexneri*. The ΔTn*10* observed in this study has the highest similarity to regions from *Enterobacter hormaechei* IncHI2 plasmids (>99% nucleotide identity and 97% query coverage). Furthermore, the ΔTn*9* (24) examined in this study was BLASTn-detected (with >99% nucleotide identity and 100% query coverage) in *Klebsiella pneumoniae* IncF plasmids.

Although *adhC* genes facilitating resistance to formaldehyde which are native to *Acinetobacter* have been reported (25), we have identified locus, likely originating from an *Enterobacter hormaechei* IncHI2 plasmid, or a *Klebsiella pneumoniae* (or *quasipneumoniae*) IncF plasmid (95% nucleotide identity and 100% coverage). However, the origin of the chlorite dismutase gene remains undeciphered, while the BLASTn of the chlorite dismutase gene observed in this study results in identical hits (>96% nucleotide identity and 98% query coverage) in various species, such as *P. aeruginosa* (chromosome), *S. enterica* (IncHI2 plasmid), *K. pneumoniae* (IncF plasmid), *E. hormaechei* (chromosome), but also *Vibrio rumoiensis* (plasmid) *Raoultella ornithinolytica* (IncF plasmid), and *Leclercia adecarboxylata* (IncHI2 plasmid).

The total of 61/159 of genomes were identified to encode the same genetic cargo (ΔTn*9,* ΔTn*10*, chlorite dismutase and the formaldehyde resistance cluster) as the one present in 3 strains of ours (0788, 0482, and 0604) in the region where the T6SS locus was expected. For the rest of genomes (98/157; as 2 genomes carried the T6SS locus) not encoding T6SS machinery, we could not identify the underlying mechanisms of loss due to draft-genome assemblies and possible crucial information missing. However, the absence of genetic cargo expected to replace the T6SS locus could suggest either a different way of T6SS locus displacement or further genomic rearrangement, possibly leading to formation of further subclades of the *A. baumannii* ST19 lineage. Alternatively, the subclade with no detectable T6SS locus displacement perhaps never encoded the T6SS in the first place.

From the total of 1287 genomes, 179 different sequence types of *A. baumannii* lacking the T6SS were detected. Furthermore, 35 genomes were typed as novel ST, with at least one novel allele, and therefore some ST might be repetitive. While additional non-ST19 *A. baumannii* lineages were detected that lack the T6SS, the specific displacement and acquisition of the ΔTn*9*, ΔTn*10*, *frmRAB*, and chlorite dismutase gene combination was observed only in *A. baumannii* ST19. The lineages *A. baumannii* ST19 (except for two genomes) and ST94 (SLV of ST19) globally lacked the T6SS. While the most genomes lacking the T6SS were detected from *A. baumannii* ST2 lineage, this is skewed by the overrepresentation of this lineage in the dataset from BIGSdb (n=6402) and the T6SS might have been replaced or lost, as observed in other cases of clinical isolates lacking some or all T6SS genes (35, 36).

The *A. baumannii* lineage ST94 (SLV of ST19) completely lacked the T6SS, however no *frmRAB* or chlorite dismutase genes were detected in this lineage. Unfortunately, the lack of complete chromosomal sequences of *A. baumannii* ST94 lineage from this dataset hinders resolution of the nature of the T6SS loss.

The loss of the T6SS in entire lineages of *A. baumannii* could also be used as an epidemiological marker. While there were reports of sporadic loss of part or the entire T6SS locus in *A. baumannii* clinical isolates (37, 38), the lineage-wide loss could be an indicator for the detection of a difficult-to-treat lineage, widely resistant to disinfectants and antibiotics, with high persistence potential.

### Conclusion

*A. baumannii* ST19 has been in occurring in multiple spikes in prevalence throughout last two decades, especially correlating with war conflicts such as the conflict in Iraq and in Ukraine. The genetic background of this lineage suggests its successful adaptation to the hospital environment. The loss of T6SS in the (nearly) entire lineage of *A. baumannii* ST19 points towards a beneficial evolutionary step, a fitness trade-off towards favouring environments where antibiotic resistance is key for survival. The increasing prevalence of the
*A. baumannii* ST19 lineage and its antibiotic and non-antibiotic (environmental stresses and disinfectants) resistance is highly concerning and warrants greater attention in surveillance efforts.

## Material and Methods

### Data Summary

The whole genome sequencing data were deposited in GenBank under following accession numbers; strain 0788 - SAMN46706486, strain 0482 - SAMN48114201 and strain 0604 - SAMN46706488.

### Bacterial isolates, genomes and metadata

Three isolates of *A. baumannii* ST19 were obtained from two different patients (patient 1 - wound, rectal swab; patient 2 - rectal swab). A further 156 genomes (including 97 from Luo et al. (17)) of *A. baumannii* ST19 were retrieved from public repositories (GenBank and BIGSdb) as of November 2025. In order to analyse other lineages of *A. baumnannii* for T6SS locus deletion, all available genomes o *A. baumannii* in BIGSdb (n=8218) were downloaded and screened.

### DNA extraction and whole genome sequencing

The DNA was extracted using DNeasy Blood & Tissue Kit (QIAGEN). A rapid barcoding kit (V14) (ONT) was then used for library preparation which was then subjected to long-read whole-genome sequencing using the R10.4.1 flowcell with the PromethION (P2 Solo) platform by Nanopore.

### Genome assembly and annotation

The long-read sequences were basecalled using the SUP model and adaptor and quality (Q ≤ 15) trimmed using Porechop v0.2.2 (https://github.com/rrwick/Porechop) and NanoFilt v2.8.0 (26), respectively. The reads were then assembled using Flye2.9.3 (27).

### Phylogenetic analysis

The maximum-likelihood trees were constructed by Prokka (28) generated gff files used for the core-genome alignment created by Roary (29). RAxML (30) was used for the calculation of the phylogenetic tree under GTRGAMMA model of rate heterogeneity method supported by 500 bootstraps. The tree was visualized in iTOL (31).

### Identification of resistance determinants

The resistance and virulence genes were detected using ABRicate (https://github.com/tseemann/abricate) employing ResFinder 4.0 (32) and VFDB 2025 (33) databases, respectively, with a 90% threshold for both gene identity and coverage. The typing of capsule-encoding loci (KL) and lipooligosaccharide outer core loci (OCL) was determined using Kaptive (34). The ABRicate (https://github.com/tseemann/abricate) was used to detect the presence of plasmid replicons using AcinetobacrPlasmidTyping database (https://github.com/MehradHamidian/AcinetobacterPlasmidTyping) (21), while the genes encoding GIs*ul2*, chlorite dismutase, formaldehyde resistance genes, and T6SS-encoding genes were detected using a custom database.

## Ethical statement

The clinical isolates sequenced in this study were obtained within retrospective study conducted under the ethical approval granted by the C. H. U. Brugmann CE 2023/51.

## Acknowledgements

AV is a recipient of a junior postdoctoral fellowship of the Research Foundation – Flanders (file number 1287223N). CVDH is supported by the Flanders Institute for Biotechnology (VIB). KN was funded by the European Union under Horizon Europe: MSCA Actions (Project 101105027) and by a junior postdoctoral fellowship from the Research Foundation - Flanders (file number 1251624N). Sequencing was performed by VIB Nucleomics Core (www.nucleomics.be).

---

**Supplementary Figure 1:**
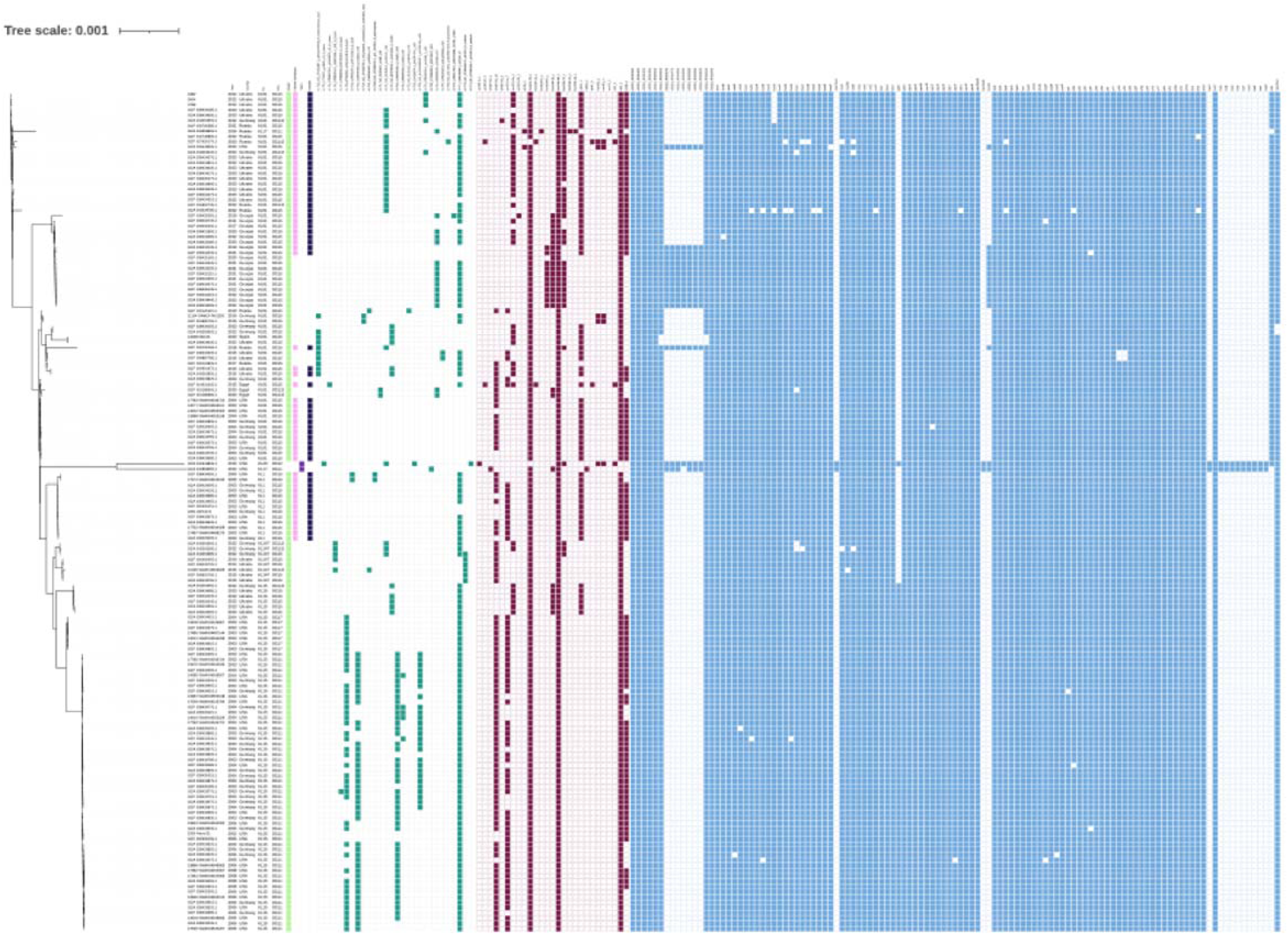
Phylogenetic tree of all *A. baumannii* genomes available in BIGSdb that do not encode the T6SS. The cluster containing the ST19 lineage is highlighted; turquoise indicates ST19, and gray indicates non-ST19.

**Supplementary Figure 2:**
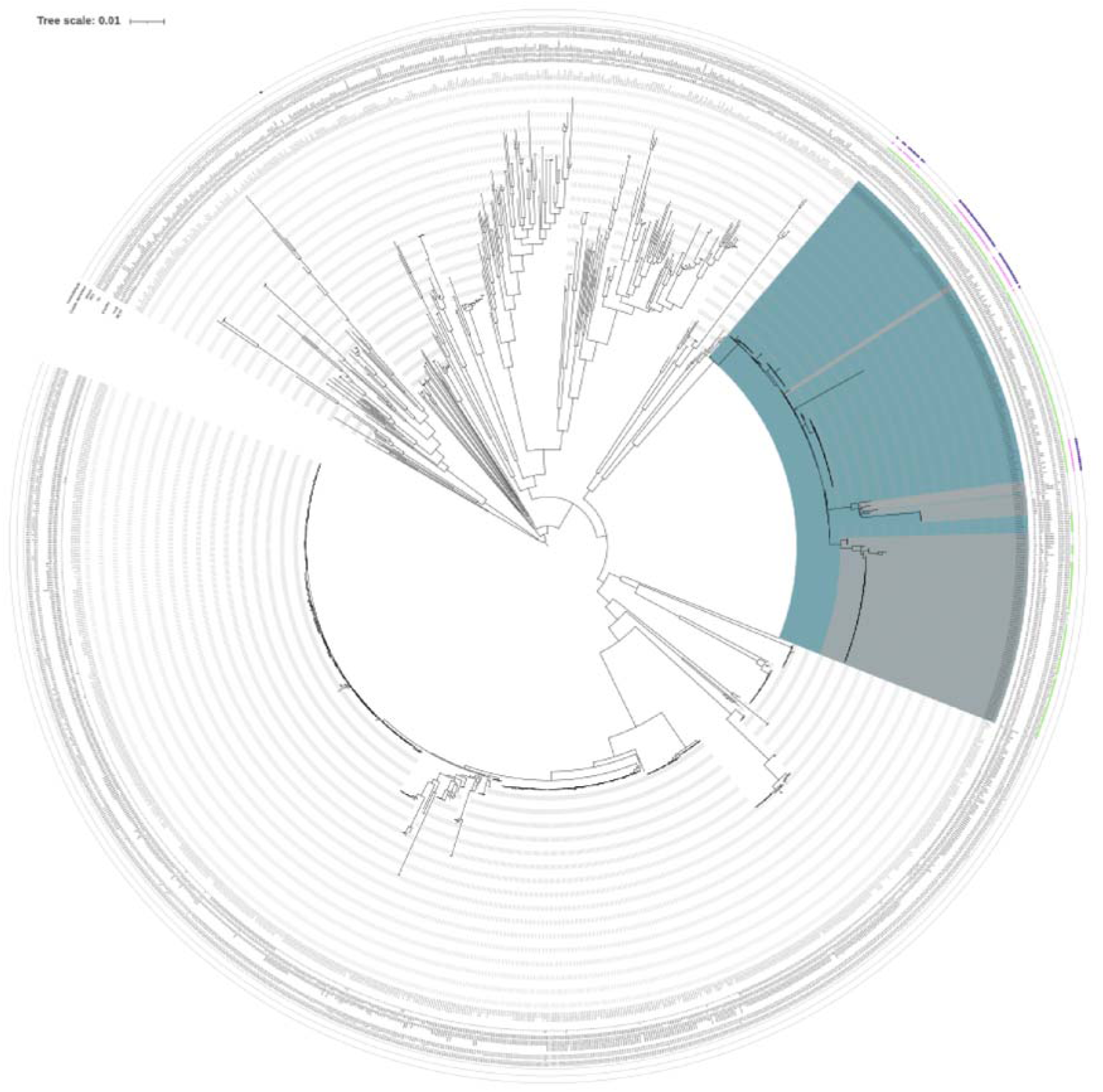
Phylogenetic analysis of three isolates and an additional 156 *A. baumannii* ST19 genomes, depicting the year of isolation, country, capsule locus type (KL), outer core lipooligosaccharide type (OCL), and the presence of the GI*sul2* genomic island, chlorite dismutase, T6SS, the *frmRAB* operon, plasmid replicons, and resistance and virulence genes, respectively.

**Supplementary Figure 3:**
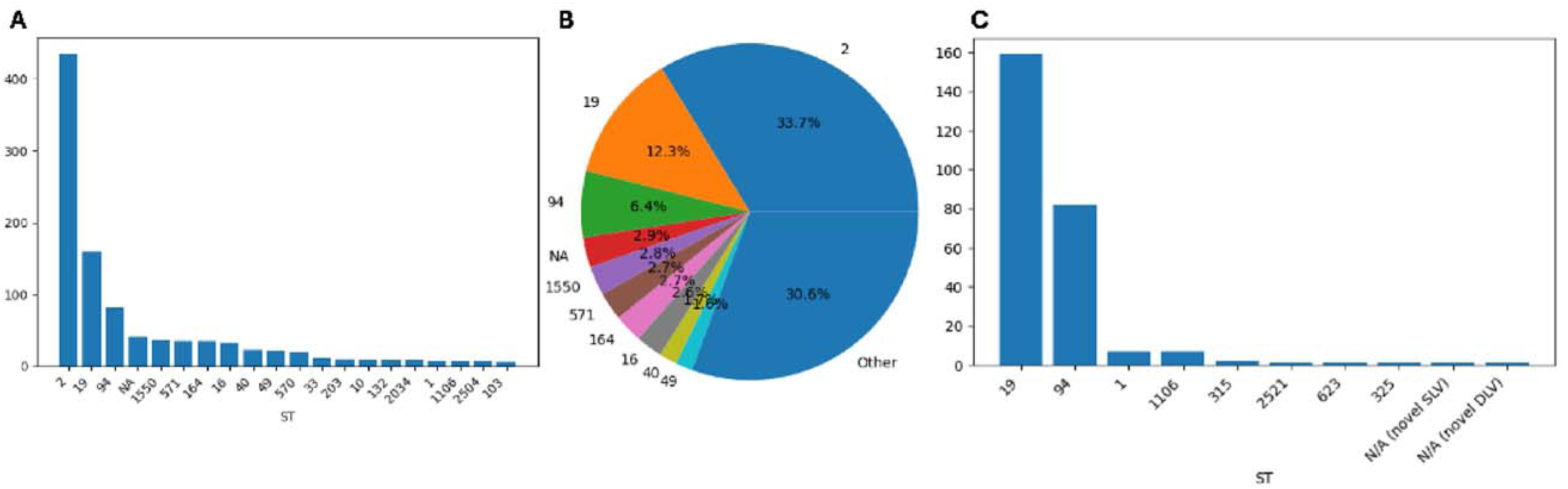
A, Quantification of the 20 most prevalent sequence types (STs) of *A. baumannii* genomes from BIGSdb lacking the T6SS. B, Pie chart showing the percentage distribution of the 10 most prevalent STs of *A. baumannii* genomes from BIGSdb lacking the T6SS. C, Numeric prevalence of STs within the cluster containing the ST19 lineage shown in Supplementary Figure 1.

